# An Integrated Workflow for Enhanced Taxonomic and Functional Coverage of the Mouse Faecal Metaproteome

**DOI:** 10.1101/2020.11.17.386938

**Authors:** Nicolas Nalpas, Lesley Hoyles, Viktoria Anselm, Tariq Ganief, Laura Martinez-Gili, Cristina Grau, Irina Droste-Borel, Laetitia Davidovic, Xavier Altafaj, Marc-Emmanuel Dumas, Boris Macek

**Author notes:** To whom correspondence should be addressed: Prof. Dr. Boris Macek Proteome Center Tuebingen Interfaculty Institute for Cell Biology Auf der Morgenstelle 15 72076 Tuebingen Germany Phone: +49/(0)7071/29-70558 Fax: +49/(0)7071/29-5779.

## Abstract

Intestinal microbiota plays a key role in shaping host homeostasis by regulating metabolism, immune responses and behaviour. Its dysregulation has been associated with metabolic, immune and neuropsychiatric disorders and is accompanied by changes in bacterial metabolic regulation. Although proteomics is well suited for analysis of individual microbes, metaproteomics of faecal samples is challenging due to the physical structure of the sample, presence of contaminating host proteins and coexistence of hundreds of microorganisms. Furthermore, there is a lack of consensus regarding preparation of faecal samples, as well as downstream bioinformatic analyses following metaproteomics data acquisition. Here we assess sample preparation and data analysis strategies applied to mouse faeces in a typical mass spectrometry-based metaproteomic experiment. We show that subtle changes in sample preparation protocols may influence interpretation of biological findings. Two-step database search strategies led to significant underestimation of false positive protein identifications. Unipept software provided the highest sensitivity and specificity in taxonomic annotation of the identified peptides of unknown origin. Comparison of matching metaproteome and metagenome data revealed a positive correlation between protein and gene abundances. Notably, nearly all functional categories of detected protein groups were differentially abundant in the metaproteome compared to what would be expected from the metagenome, highlighting the need to perform metaproteomics when studying complex microbiome samples.

## Introduction

The prokaryotic component of the gut microbiota has multiple roles, contributing to carbohydrate fermentation and maintenance of gut barrier integrity, as well as antimicrobial and immunomodulation activities [1, 2]. In metabolically healthy humans and mice, the gut microbiota is predominated by two to three bacterial enterotypes [3–5]. These enterotypes display significant heterogeneity in terms of species number, composition and relative abundances depending on the location of the sample (upper vs lower gastroinstestinal tract) or the timing (circadian variations) [6, 7]. The gut microbiota has recently been associated with conditions ranging from inflammatory bowel syndrome to Parkinson’s disease [8–11]. An increasing number of studies have reported associations between the gut microbiota and neurodevelopmental disorders [12–14]. This includes changes in the gut microbiota of Down syndrome individuals in comparison to non-trisomic individuals [15]. Given the established interaction between the host and the gut microbiota, a functional analysis of the gut microbiome may help in understanding its contribution to pathophysiology.

In this context, approaches relying on nucleotide sequencing have so far been preferred by the scientific community due to lower experimental costs, higher data throughput and proven analytical workflows. While metagenomics assesses the genetic potential, metaproteomics investigates gene products (and therefore functions). However, metagenomics usually provides more in-depth information in comparison to metaproteomics, for example due to the higher dynamic range of detection. In particular, microbiome functional analysis can be performed using high-resolution mass spectrometry (MS), to measure either protein abundance or metabolite production [16–18]. Although bacterial MS-based proteomic approaches are well established, metaproteomic sample preparation is hindered by many challenges, such as physical structure of the sample, the presence of contaminating proteins and the coexistence of hundreds of microorganisms.

Many studies have reported increased protein identification due to laboratory optimisation for the analysis of metaproteome samples [19–23]. In humans, different sample preparation methodologies have been shown to result in significant changes in the taxonomic composition and functional activities representated [19,24,25]. Beyond sample preparation, the bioinformatic processing of metaproteomic data remains challenging, due to the choice of representative protein sequence database, elevated false discovery rate for peptide identification and the redundancy in protein functional annotation. Some of these challenges have already been addressed by published software packages, such as MetaProteomeAnalyzer [26] and MetaLab [27], which are all-in-one metaproteomic analytical workflows, or UniPept [28], which allows peptide-based taxonomic representation. In addition, the choice of protein sequence database has been shown to play a major role in protein identification from metaproteome samples, with notably matching metagenome-derived protein sequence databases displaying the best identification rate performance [29–32]. Previous studies have also investigated ways to determine taxonomic representation from metaproteome samples, which has been shown to differ between metagenome (bacterial presence) and metaproteome (bacterial activity) [32, 33].

Here, we present a state-of-the-art MS-based workflow for the optimal metaproteome characterisation of murine faecal samples. We focused on a number of aspects that remain under-investigated in murine stool samples: (1) the impact of sample preparation methods, namely low speed centrifugation (LSC) and no LSC (nLSC), on protein identification and taxonomic representation; (2) the high false positive rates in searches involving very large databases; (3) the differences in taxonomic annotation of MS-identified peptides based on different software; and (4) the lack of assessment of the functional enrichment provided by the metaproteome compared to its matching metagenome potential.

## Results

### Low-speed centrifugation increases peptide identification rates

Our initial experiment involved the establishment of an optimal sample preparation workflow applied to the mouse faecal metaproteome. In this context, we assessed two critical steps within our sample preparation method: 1) the usage of LSC; and 2) in-solution digestion versus filter- aided sample preparation (FASP) (**Figure S1A**, **Table S1**).

**Table 1:** Performance comparison of different sample preparation and data analysis steps. In bold are the best methods according to assessed criteria: peptide/protein count, host/dietary contamination, *Firmicutes* or *Bacteroidetes* representation, time efficiency, FDR, identification rate, taxon-assigned peptides and number of taxonomic identification precision. The performance status is displayed using minus sign for poor, equal sign for similar/no difference or plus sign for good performance.

|  |  | Peptide/protein count | Host/dietary contamination | Firmicutes | Bacteroidetes | Time efficiency | FDR | Identification rate | Taxon assigned peptides | Precision |
| --- | --- | --- | --- | --- | --- | --- | --- | --- | --- | --- |
| Centrifugation | **LSC** | + | - | - | + | - |  |  |  |  |
|  | nLSC | - | + | + | - | + |  |  |  |  |
| Digestion | **In-solution** | + |  |  |  | - |  |  |  |  |
|  | FASP | - |  |  |  | + |  |  |  |  |
| Search strategy | **Single-step** | - |  |  |  | + | + | - |  |  |
|  | Two-step protein | + |  |  |  | - | -- | + |  |  |
|  | Two-step sections | + |  |  |  | -- | - | + |  |  |
|  | Two-step taxa | - |  |  |  | - | + | - |  |  |
| Taxon quantification | Kraken2 |  |  |  |  | + |  |  | + | -- |
|  | Diamond |  |  |  |  | -- |  |  | - | - |
|  | **Unipept** |  |  |  |  | + |  |  | + | + |

The number of peptide spectral match (PSM) identified per MS raw file in the LSC group was significantly higher with 26 % more identifications (**Figure 1A**). This was also observed at the peptide and protein group level, but to a lower extent for the latter. Approximately 15 % of protein groups were identified by a single peptide, while the median protein sequence coverage was 18.7 %. Such metrics are usually indicative of highly complex samples that are not completely covered by a single MS measurement under the stated parameters.

**Figure 1:**
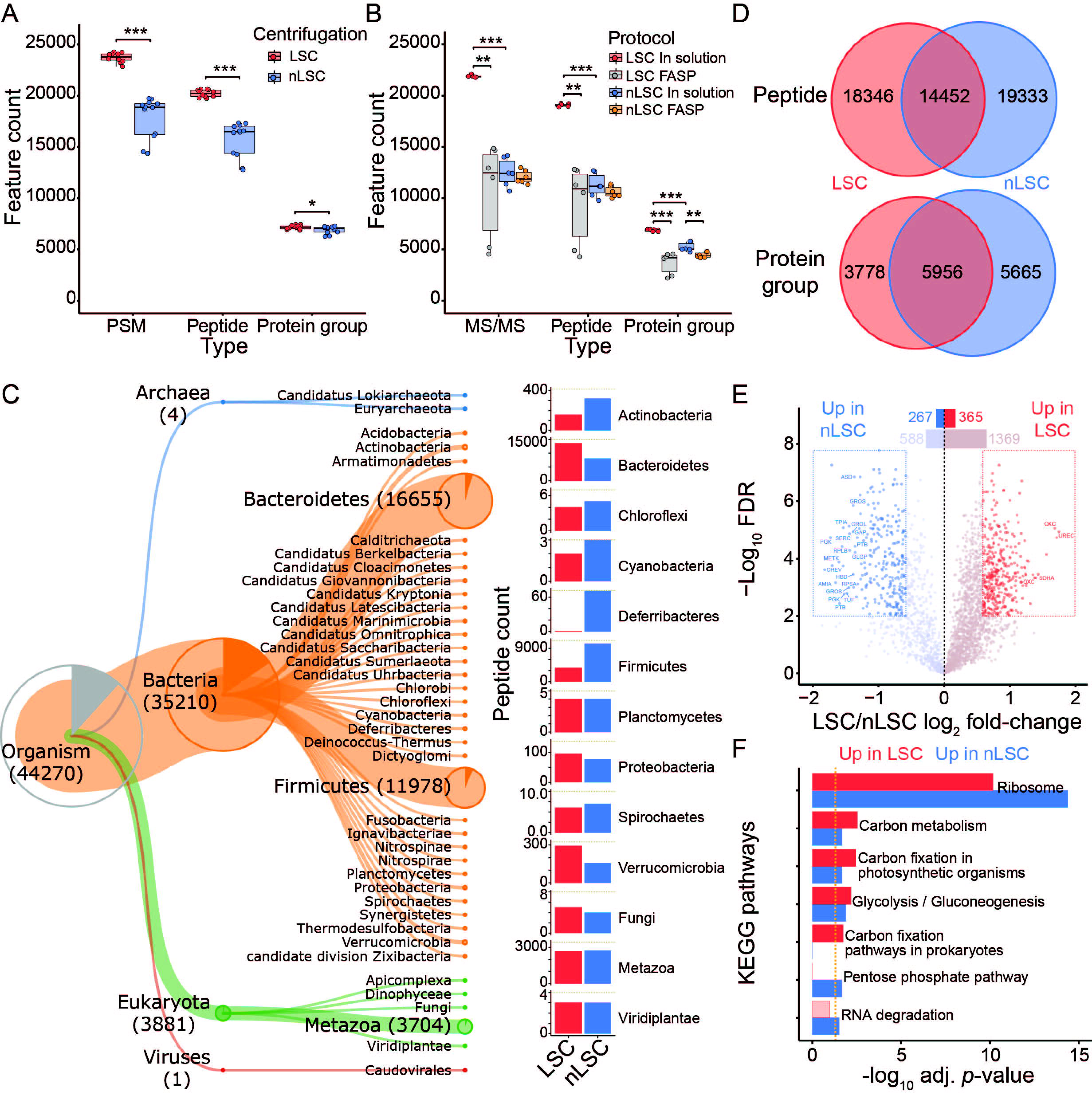
Low speed centrifugation impacts protein identification and taxonomic representation. A) Number of MS/MS spectra, peptides and protein groups per samples for the comparison between LSC (red) and nLSC (blue) methods. B) Number of identified MS/MS spectra, peptides and protein groups per samples for the comparison between LSC-in solution digestion (red), LSC-FASP (grey), nLSC-in solution digestion (blue) and nLSC-FASP (orange) methods. A-B) Represented significance results correspond to t-test on N = 12 (A) or N = 6 (B): * *p*- value ≤ 0.05, ** ≤ 0.01, *** ≤ 0.001. C) Hierarchical representation of Unipept- derived taxonomy (down to phylum level) for the peptide identified in the LSC and nLSC. The barplot represent the taxonomic abundance for LSC (red) and nLSC (blue) methods based on peptide counts (only for taxon identified with 3 or more peptides). D) Overlap in the overall identified peptides or protein groups between the LSC and nLSC methods. (E) Volcano plot of the protein abundance comparison between LSC and nLSC approaches. Significant protein groups based on paired t-test from N = 12 with FDR ≤ 0.01 and absolute fold-change ≥ 2.5. (F) KEGG pathways over-representation testing for the protein groups that significantly increase (red) or decrease (blue) in abundance between LSC and nLSC sample preparation approaches. Fisher exact-test threshold (gold dotted line) set to adjusted *p*-value ≤ 0.05.

In-solution digestion consistently outperformed FASP based on PSMs, peptides and protein groups identification (**Figure 1B**). Compared to other methods, in-solution digestion combined with the LSC procedure provided nearly twice as many PSM or peptide identifications and 30 % more protein groups. Furthermore, there was much less variability in the number of peptides and protein groups identified across samples with this method.

### LSC aids in recovery of *Bacteroidetes* proteins, whereas nLSC favours *Firmicutes* and *Deferribacteres* proteins

Peptides identified after LSC and nLSC were analysed to identify their phylogenetic origin. The lowest common ancestor was determined using the Unipept interface [34], which assigns peptide sequences to taxa. The most abundant superkingdom consisted of bacteria, among which two taxa were highly represented in both LSC and nLSC, namely *Bacteroidetes* and *Firmicutes* (**Figure 1C**, **Table S1**). However, there were large differences in the number of peptides assigned to these two main bacterial phyla when comparing LSC and nLSC methods. *Bacteroidetes* accounted for 66 % and 37 % of peptides, whereas *Firmicutes* amounted to 18 % and 47 % of peptides in LSC and nLSC procedures, respectively. In addition, *Actinobacteria* and *Deferribacteres* showed a higher taxonomic representation in nLSC compared to LSC, whereas *Verrucomicrobia* showed an opposite trend.

Based on peptides identification, Eukaryota was the second most abundant superkingdom and consisted mostly of metazoan hits. Under the assumption that these eukaryotic peptide sequences originated from the host, the proportion of *Mus musculus* proteins was investigated further using intensity-based absolute quantification (iBAQ) values. The LSC samples contained on average nearly two-fold more murine proteins (20.4 %) in comparison to nLSC samples (14.6 %) (**Figure S1B**). Such findings were surprising since the use of the LSC method was reported in a previous study to help with the removal of human cells [24]. We also investigated the presence of peptides from host diet and found very low levels of dietary peptides contamination (approximately 2 %), which was higher among LSC-prepared samples (**Figure S1C**). As previously reported, we show that the majority of dietary proteins are absent or depleted during the initial solubilisation step of the faecal pellet, a step common to both procedures [35]. Overall, our results show that LSC and nLSC methods favour the recovery of different taxa, suggesting that both methods have merits and may be used in combination.

### LSC and nLSC methods are characterised by different protein abundance profiles

We further investigated the overlap between the peptides or protein groups identified following either LSC and nLSC procedures (**Figure 1D**). In terms of peptides, only 27.7 % were identified with both procedures, the rest of the peptides being split equally into unique to LSC and nLSC methods. Similar results were observed at the protein groups level with 38.7 % of protein groups being identified in both procedures. This was illustrated further through a principal component analysis (PCA), showing separation of samples based on centrifugation methods, as well as clustering of technical replicates (from cell lysis step) (**Figure S1D**). Label-free quantitative (LFQ) comparison between LSC and nLSC procedures revealed an intermediate correlation (ρ = 0.44) (**Figure S1E** and **F**). Besides, LFQ correlation among the samples prepared via LSC was superior to samples prepared with nLSC (**Figure S1G**). Our findings indicate that while the two procedures have a poor identification overlap, the main differences may still result from biological variations.

Using LFQ intensities, we then performed a *t*-test to identify which protein groups have different abundances between the two procedures. Out of 2,589 quantified protein groups, 365 and 267 showed a significant increase and decrease in abundance between LSC and nLSC samples, respectively (FDR ≤ 0.01 and absolute fold-change ≥ 2.5) (**Figure 1E**, **Table S1**). We gained functional insights into these differences by performing an over-representation analysis of KEGG pathways. The over-represented pathways based on the up- or down-regulated protein groups were mostly similar (FDR ≤ 0.05) and were associated with core microbial functions, such as ribosome, carbon metabolism and carbon fixation pathways (**Figure 1F**, **Table S1**). The protein groups unique to LSC or nLSC showed over-representation of protein export in the LSC samples, whereas biosynthesis of amino acid, fatty acid degradation and bacterial chemotaxis were over-represented in the nLSC samples (**Figure S1H**). Protein differential abundance testing confirmed the divergence between LSC and nLSC procedures and was suggestive of broad taxonomic changes, rather than variation in functional activities.

### Two-step database search strategy shows a dramatic increase in false positive rate

After measurement via liquid chromatography coupled to tandem mass spectrometry (LC-MS/MS) and acquisition of LC-MS/MS raw data, the MS/MS spectra are searched against a protein sequence database. One aspect of database search is the controversial use of a two-step search strategy, whereby LC-MS/MS measurements are processed initially against a large protein sequence database with no FDR control (FDR ≤ 1%). Subsequently, the original database is filtered to retain only protein sequences that were identified during the first search. During the second database search, the measurements are processed against the reduced database with FDR control (e.g. FDR ≤ 0.01) [36]. To assess these search strategies, we searched a single HeLa cell LC-MS/MS file using MaxQuant software against a *Homo sapiens* protein sequence database supplemented with different number of bacterial protein sequences (**Figure S2A**). The HeLa measurement is used here as a proxy for a complex microbiome measurement, with the exception that the sample composition is known and from a single organism.

We initially established a benchmarked standard by processing the HeLa measurement only against an *H. sapiens* database, which resulted in approximately 5,000 human (eukaryota) protein groups identified for the single-step search at FDR ≤ 0.01 (**Figure 2A**, **Table S2**). Notably, the same database used in a two-step search identified less than 1 % additional protein groups in comparison to a single-step search, despite nearly twice as much processing time. We then processed our HeLa measurement against the *H. sapiens* database supplemented with 1×, 2×, 5×, 10× and 20× bacterial protein sequences, resulting in increasingly large databases (**Figure S2A**, **Table S2**). For the single-step database search against the 1:20 database, we observed a 10 % decline in the number of human protein groups identified, while 132 bacterial protein groups were identified (false positives). On the contrary, the 1:20 two-step database search resulted only in a 1 % decrease compared to the benchmarked standard. This processing also revealed a large number of bacterial protein groups identification (980 protein groups). Furthermore, the two-step search led to large number of MS/MS spectra to be assigned to different sequences (or newly assigned) in comparison to the benchmarked standard (**Figure S2B**, **Table S2**); this phenomenon was much less pronounced when performing the single-step search.

**Figure 2:**
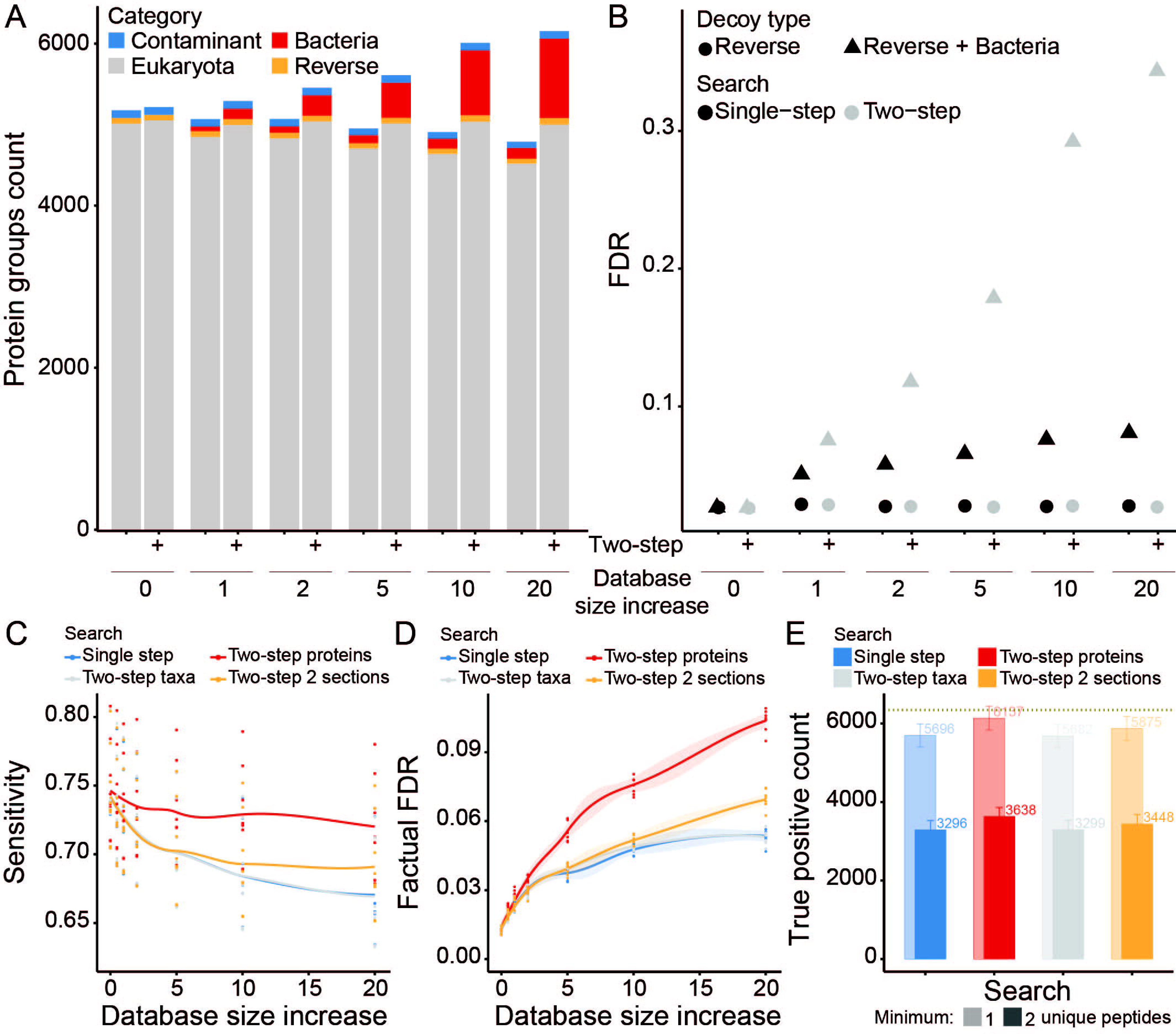
Two-step database search in combination with target-decoy strategy leads to a dramatic increase in false positive rate. A) The protein groups count is shown for single- or two-step search strategies across increasingly large protein sequence databases. Counts are colour-coded per category, with eukaryote (grey), bacteria (red), contaminant (blue) and reverse (orange) hits. B) The FDR is calculated for single- or two-step search strategies across increasingly large protein sequence databases. The FDR is calculated based on reverse hits only (circle shape) or reverse plus bacterial hits (triangle shape). C & D) The sensitivity (C) and factual FDR (D) based on protein groups identification across increasingly large protein sequence databases. The compared database search strategies are single-step (blue), two-step taxon filtering (grey) and two-step protein filtering without (red) or with (orange) database sectioning. Lines represent the median (and the shading corresponds to the standard error) from N = 8 LC-MS/MS runs. E) The true positive count based on protein groups identified with a minimum of one (shaded colouring) or two (unshaded colouring) unique peptides for the largest database (i.e. 20). The compared database search strategies are single-step (blue), two-step taxon filtering (grey) and two-step protein filtering without (red) or with (orange) database sectioning. Bars and numbers indicate the median count, while error bars correspond to the standard deviation, from N = 8 LC-MS/MS runs. The overall maxima of true positive count based on single-step search is indicated as a horizontal dotted line (gold).

We then calculated the factual FDR for each processing approach using either the reverse hits or the reverse hits plus the bacterial hits (which in our case are false positives). For both the single-step and the two-step search, we obtained an FDR of 2.6 % when using only the reverse hits for FDR calculation (**Figure 2B**). However, when using the reverse hits plus the bacterial hits, we calculated a factual FDR of 8 % and 34 % for the single- and two-step search with 1:20 database, respectively. This represents a dramatic increase in the rate of false positive identification when using two-step search, despite controlling for 1 % FDR. Notably, these false positive hits would remain unnoticed in a microbiome sample of unknown composition, thus highlighting the inherent problem associated with the two-step database search.

### Optimisations of the two-step database search cancels out its higher sensitivity

To further assess database search strategies in a metaproteomic context, we retrieved a metaproteome dataset of known taxonomic composition that was published by Kleiner and colleagues [32]. This dataset consisted of 32 organisms of uneven abundances, including bacteria (25), archaea (1), eukaryotes (1) and viruses (5). We processed eight LC-MS/MS measurements against a database containing the proteomes of these 32 organisms, which we supplemented with 0.5×, 1×, 2×, 5×, 10× and 20× bacterial protein sequences, resulting in increasingly large databases. We then compared the results obtained from single-step search strategy against: (1) “two-step protein” search to keep identified proteins [36]; (2) “two-step taxa” search to keep identified taxa [30]; and (3) “two-step two sections” search to keep identified proteins after sectioned search [37]. While all search strategies resulted in similar accuracies, the “two-step protein” search maintained a high sensitivity even when using large databases (i.e. 20×) (**Figure 2C** and **S2C**). However, upon investigation of the factual FDR (reverse hits plus the false bacterial hits), the “two-step protein” search resulted in twice as many false positive identifications compared to the single-step search (**Figure 2D**, **Table S2**). Similar results were also observed when focusing on the precision (**Figure S2D**). Our investigations revealed that the “two-step taxa” search behaved nearly identically to the single-step search, whereas the “two-step two sections” search displayed performance in-between the first-step and “two-step protein” searches.

Because, all assessed search strategies underestimated the real FDR, we attempted to identify any particularity of the false positive protein groups identification and thus focused on processings against the largest database (20×). We show that the median number of unique peptides (i.e. peptides that are uniquely assigned to a protein group) are 1 and 2 for the false and true positive hits, respectively (**Figure S2E**). We then compared results obtained using a post-processing filtering step requiring a minimum of 1 or 2 unique peptides per protein groups. Our results show that requiring a minimum of 2 unique peptides would efficiently control the FDR (≤ 1%) at the expense of a significant drop in protein identification (**Figure 2E** and **S2F**, **Table S2**). This investigation of different database search strategies applied to metaproteome samples further highlighted the limitations (i.e. factual FDR) of two-step searches, even following optimisation (i.e. sectioned search) or filtering.

### Unipept software provides the most accurate and precise taxonomic annotation

Another important aspect of metaproteomic studies is the determination of taxonomic activity (protein biomass), which has been reported to differ from taxonomic representation derived from metagenomic studies [32]. While it is straightforward to compute taxonomic activity from the abundance of peptides (or proteins) of known taxonomic origin, there has not been an exhaustive assessment of software that can taxonomically annotate MS-identified peptides. Here, we assessed three software—i.e. Kraken2 [38], Diamond [39] and Unipept [28]—with regards to their taxonomic annotation performance on the dataset from Kleiner and colleagues [32]. The Kraken2 software provided consistently higher percentage of peptides that could be taxonomically annotated (c.a. 18% peptides annotated to species level), followed by Unipept (5%) and Diamond (1%) (**Figure 3A**). However, Kraken2 also identified a very large number of taxa that were not present in the artificial samples from Kleiner and colleagues (**Table S3**) and thus would be false positive hits. Unsurprisingly, these false positive hits were characterised by low PSM counts in comparison to true positives (**Figure S3A**). This led us to assess these software packages in terms of accuracy, precision, sensitivity, specificity and F- measure for taxonomic identification using a range of PSM count thresholds (**Figure 3B** and **S3B-E**). In this context, the Unipept software significantly outperformed Kraken2 and Diamond, especially with regard to the F-measure and precision. Notably, the implementation of a minimum PSM count threshold (i.e. between 1 and 5) resulted in accuracy, precision and specificity improvements for all software, but at the cost of a reduced sensitivity.

**Figure 3:**
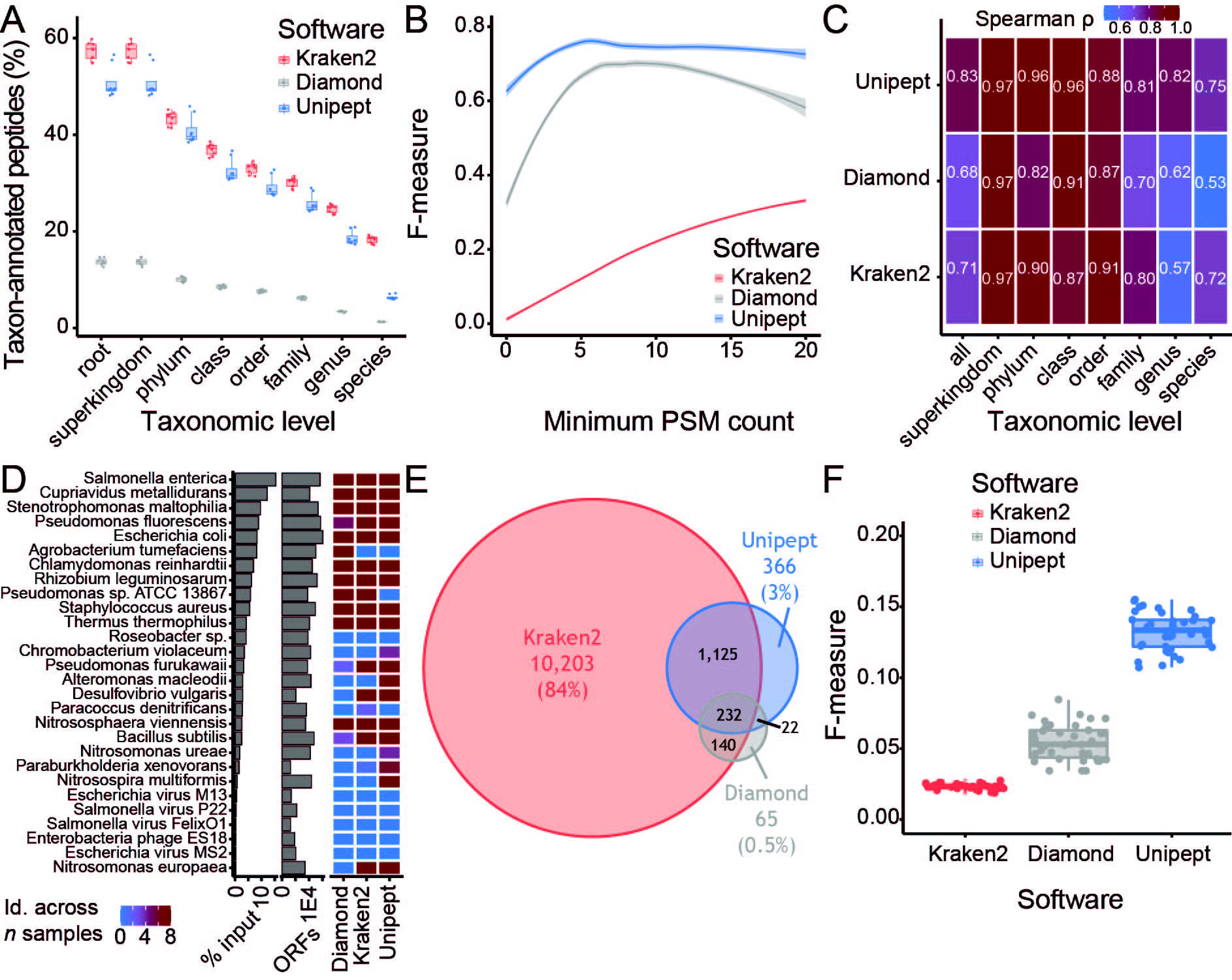
Unipept software provides the most precise taxonomic annotation of MS-based peptide identification. A) Percentage of taxon-annotated peptides at each taxonomic level for the comparison between Kraken2 (red), Diamond (grey) and Unipept (blue) software. B) Assessment of the impact of the minimum number of PSM count per taxon onto the F-measure for taxonomic annotation. The F-measure was compared between Kraken2 (red), Diamond (grey) and Unipept (blue) software. C) Heatmap representing the correlation (Spearman ρ) in taxonomic abundance between sample input protein (expectation) and different taxonomic annotation software (i.e. Kraken2, Diamond and Unipept). The correlation was performed overall, as well as for each taxonomic level. D) Organisms pooled in artificial samples are ranked based on the protein material input, as displayed in the left-most barplots (x-axis in log10 scale). The proteome size (ORFs) for these organisms on UniProt web resource is displayed in the right-most barplot (x-axis in log10 scale). The heatmap compares the taxon identification across samples between Kraken2, Diamond and Unipept. A-D) Samples from the study by Kleiner and colleagues, with N = 8. E) Overlap in the overall identified taxa between the Kraken2 (red), Diamond (grey) and Unipept (blue) software. F) A comparison of the F-measure distribution for taxonomic annotation between the Kraken2 (red), Diamond (grey) and Unipept (blue) software. Each point represents an individual mouse. E-F) Samples from this study using mouse faecal material, with N = 38.

Thus, without a PSM count threshold, we correlated the taxonomic abundance derived from each software annotation against the known input protein from Kleiner and colleagues’ artificial samples (**Figure 3C**). Overall, the Unipept software provided the highest correlation (Spearman ρ = 0.83), as well as at most taxonomic levels (including species). Interestingly, the dynamic range of taxon detection by MS spanned two orders of magnitude, with *Salmonella enterica* being approximately 230 times more abundant than *Nitrosomonas europaeae* (**Figure 3D**, **Table S3**). Unipept was also the only software allowing identification of *Nitrosomonas ureae*, *Paraburkholderia xenovorans* and *Nitrosospira multiformis*. Importantly, none of the software could identify the five viral organisms present in the samples, the reason being technical since no peptide coming from those viral proteins was detected by MS. Finally, we assessed the impact of different database search strategies on taxonomic abundance derived by the Unipept software (**Figure S3F**). Similarly to our findings from the previous section, the F- measure metric highlighted the superiority of single-step strategy when it comes to taxonomic identification. Taken together, we show that, based on different metrics and samples of known composition, the Unipept software provides better taxonomic annotation in comparison to Kraken2 and Diamond.

### The complex microbial composition of faecal samples is best recapitulated by the Unipept software

To check whether our results are also applicable to the microbial composition of faecal samples, we prepared samples using the LSC method from faeces collected in a cohort of 38 mice. The resulting LC-MS/MS data were processed using a single-step search strategy against a matching metagenome protein database (with no knowledge of taxonomic composition). We initially annotated the MS-identified peptides using Kraken2, Diamond and Unipept, which revealed an overlap of 232 taxon (1.9%) between all three software. Such low overlap was largely driven by the suspected large number of false positive hits identified by Kraken2 (10,203 uniquely identified taxon), as seen in the previous section. We then performed pairwise correlation between every samples combination within each software using taxonomic abundance (**Figure S3G**). While Diamond displayed a higher correlation (median spearman ρ = 0.71), this is likely driven by the small number of identified taxa, most of which at the taxonomic levels closer to the root (e.g. superkingdom, phylum) and is thus a poor performance estimate.

To determine which taxa are likely true or false positive hits, we made use of the taxonomic composition foreknowledge (at the species level only) from the mouse microbiome catalogue [40]. With this approach, the Kraken2 software showed the best sensitivity (median = 0.17) compared to Unipept (0.13) and Diamond (0.03) (**Figure S3H**, **Table S3**). Based on precision and F-measure, Kraken2 performance collapsed, whereas Unipept software had median precision and F-measure superior to 0.1 (**Figure 3F** and **S3I**). Using taxonomic foreknowledge, our findings suggest that the Unipept software provides superior predictive power for taxonomic annotation of faecal samples.

### Metaproteome to metagenome correlation highlights an over-representation in the core microbiome functions

Due to the availability of matching metagenomic and metaproteomic data for our cohort of 38 mice, we assessed the correlation between gene and protein abundances. To deal with the intrinsic differences between the two datasets, the gene entries were grouped in a similar fashion as the protein groups (i.e. based on peptide identification) and the maximum expression was calculated per gene group. Here, we show that a majority of gene-protein pairs (91 %) have a positive correlation, with a median of 0.39, the rest having a median negative correlation of 0.09 (**Figure 4A**, **Table S4**). Notably, 3,519 gene-protein pairs displayed a significant positive correlation. In addition, we compared the distribution in gene abundances depending on whether the corresponding protein was identified by MS (**Figure S4A**). As expected, it shows that MS-based proteomics only identifies a subset of proteins towards the higher abundance.

**Figure 4:**
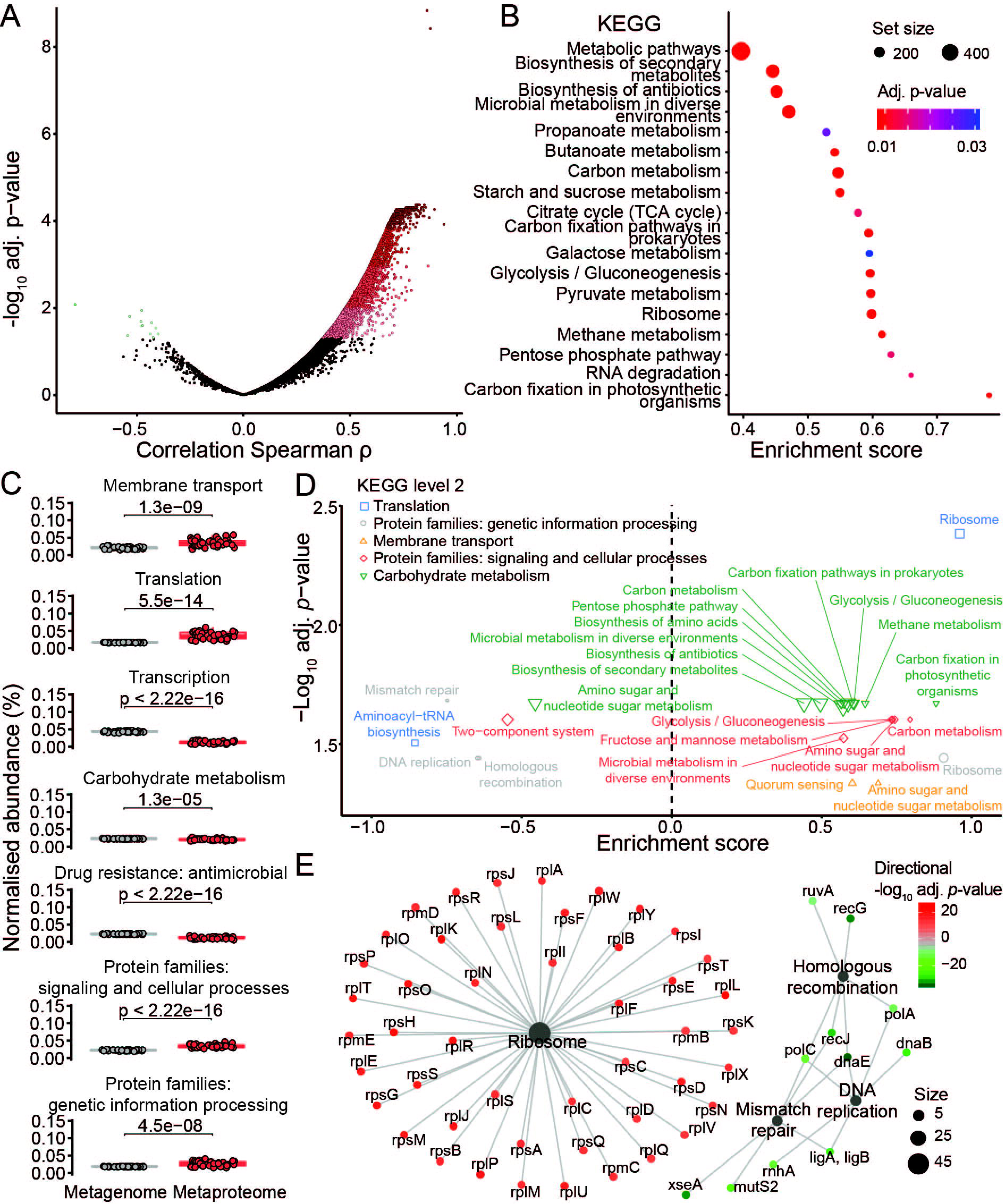
Functionally active pathways derived from the metaproteome differs from the metagenome potential. A) Correlation is shown between each protein groups (metaproteome) and corresponding gene “groups” (metagenome) abundances. Correlation was tested using Spearman’s rank correlation and p-value was adjusted for multiple testing using Benjamini-hochberg correction. Significantly correlating protein/gene groups are in red colours, while significantly anti-correlating protein/gene groups are in green colours (adjusted *p*-value ≤ 0.05). B) GSEA of KEGG pathways based on ranking of the protein/gene groups correlation. Pathway node colour corresponds to GSEA results adjusted p-value and node size matches the number of protein/gene group assigned to the pathway. C) Comparison in the proportion of selected KEGG functional categories (level 2) between metaproteome (red) and metagenome (grey). Paired t-test p-values are indicated (N = 38). D) GSEA of KEGG pathways based on ranking of t-test results from KEGG orthology proportion between metaproteome and metagenome. KEGG pathways are colour-coded based on KEGG functional categories (level 2). Only significantly over-represented KEGG pathways are shown with adjusted *p*-value ≤ 0.05. E) Interaction network between KEGG orthologies and KEGG pathways for the KEGG functional category “Protein families: genetic information processing”. Pathway node size corresponds to number of KEGG orthologies associated to it. KEGG orthologies are colour-coded based on directional adjusted *p*-value from the t-test comparison between metaproteome and metagenome.

To identify the core pathways within our mice cohort, we performed an over-representation analysis of the significantly correlated gene-protein pairs (**Figure 4B**, **Table S4**). Among these pairs, there was an over-representation in carbon fixation, glycolysis-gluconeogenesis, citrate cycle and carbon metabolism pathways (KEGG) [41]. We further characterised the correlating genes and proteins and identified 20 over-represented gene ontology molecular functions (GOMF) that were involved in ADP, ribosome, carbohydrate and electron transfer (**Figure S4B**, **Table S4**). Our results confirm the central role of carbon fixation and general metabolism, which are associated with bacterial energy production, in the murine faecal microbiome under the analysed conditions.

### The metaproteome is enriched in functionally active pathways compared to the matching potential encoded in the metagenome

The metagenome corresponds to the microbiome’s genetic potential, whereas the metaproteome represents its truly expressed functional activities. Thereby, we compared the functional abundance derived from the metagenomic versus metaproteomic datasets within our cohort of 38 mice. To allow comparison, the KEGG level 2 categories were quantified and normalised separately for each omic datasets (**Figure S4C**, **Table S4**). Out of 55 KEGG categories, we found 15 and 37 to be significantly increased and decreased in abundance at the metaproteome level in comparison to the metagenome (FDR ≤ 0.05). In general, the metagenome-based quantification of KEGG categories was stable across categories, whereas large differences were observed for the metaproteome.

To prioritise the KEGG categories, we selected eight categories differing significantly in terms of gene-protein correlation in comparison to the overall correlation (**Figure 4C** and **S4D**). Among the KEGG categories displaying higher abundance in the metaproteome compared to the metagenome were the membrane transport, translation, signalling and cellular processes, and genetic information processing. Conversely, transcription, carbohydrate metabolism and antimicrobial drug resistance exhibited lower abundance. The KEGG Orthology (KO) entries differing significantly in abundance between the metagenomes and metaproteomes were identified via *t*-test and used for gene set enrichment analysis (GSEA). GSEA revealed an enrichment of a number of overlapping KEGG pathways, with 19 and 6 pathways positively and negatively enriched, respectively (**Figure 4D**, **Table S4**). Interestingly, we found the ribosome pathway enriched in protein with increased abundance (between metaproteome and metagenome datasets), therefore highlighting the functional activation of this pathway (**Figure 4E** and **S4E**). Conversely, homologous recombination, DNA replication and mismatch repair were enriched in protein with decreased abundance, suggesting no or low activation of these pathways. Overall, our findings highlight the critical importance of metaproteomics to characterise microbiome samples particularly when it comes to their functional activity.

## Discussion

Here, we investigate some key aspects of metaproteomic workflow applied to murine faecal samples in order to enhance protein identification, taxonomic and functional coverage. We focused on the assessment of (1) different sample preparation methods, (2) strategies to control for false positive rates during database search, (3) taxonomic annotation software for accurate MS-derived taxonomic representation and (4) the importance of metaproteomics to determine functionally enriched pathways. Our results led to an overview of the strengths and weaknesses of each assessed methods (**Table 1**) in the context of murine faecal metaproteomics.

To the best of our knowledge this is one of the largest and most extensive comparisons undertaken to date, comprising over 40 different biological samples and over 50 LC-MS/MS runs. Overall, we reached identification rates that are similar to bacterial shotgun proteomics (ca. 20-40 %). In comparison to previous murine faecal metaproteomic studies, we identified more non-redundant peptides per samples (approximately 20,000 non-redundant peptides on a 60 min gradient) [42, 43]. Several parameters may have influenced such performance, among which are the use of a faster and more sensitive Orbitrap instrument (i.e. Q Exactive HF) [44, 45] and a more representative protein sequence database (i.e. mouse metagenome catalogue or mouse matching metagenome) [40]. Importantly, the impact of mass spectrometer speed and sensitivity should not be overlooked in a typical metaproteomic measurements. Indeed, the type and model of MS instrument was among the parameters with the greatest impact on identification rates. Some of our initial investigation showed significant increase in peptide and protein identification rate when using the Q Exactive HF (faster scanning, improved sensitivity) versus the Orbitrap Elite (data not shown, but downloadable from ProteomeXchange).

### Both LSC and nLSC methods have merits for the metaproteomic analysis of murine faecal samples

Our study confirms previous observation with regard to the depletion or enrichment of several major bacterial phyla, which is dependent on laboratory preparation method and specifically the usage of differential centrifugation [24]. In this context our results do not match with the study from Tanca and colleagues, who reached opposite conclusions. There are several possible explanations for these discrepancies, such as the host organism under study (*i.e. Mus musculus* versus *Homo sapiens*) and different protein sequence database construction (*i.e.* mouse microbiome catalogue versus UniProtKB custom microbiome). The LSC approach also leads to more consistent identifications and as a result fewer missing values, which is a general and extensive problem in metaproteomic datasets. Regarding the topic of reproducible protein identification and quantification, a recent metaproteomic study demonstrated the use of Tandem Mass Tag (TMT) approach in human stool samples [22].

Investigation of the differences in taxonomic composition between LSC and nLSC revealed broad changes already at the phylum level. Notably, *Bacteroidetes* and *Verrucomicrobia* were enriched within LSC-prepared samples, whereas *Firmicutes, Actinobacteria* and *Deferribacteres* phyla were over-represented in nLSC samples. Additional comparison to the phyla detected by metagenomics in the mouse microbiome catalogue study tends to agree more with the nLSC approach [40]. However, the samples from that study were also prepared using a nLSC approach, which may explain the similarity. Importantly, it has been reported that the removal of faecal particles may also lead to exclusion of proteins or organisms attached to these faecal debris [24], thus leading to a bias in the LSC approach. Our results at the protein level showed significant changes in abundance, which were indicative of broad taxonomic changes, more so than variation in functional activities. Importantly, recent studies have reported considerable changes in rodent microbiota depending on suppliers or on shipping batch, even for mice housed in identical environments [46, 47]. Murine gut microbiota is also significantly different from other mammals, such as human [40]. In this context, our results on metaproteomic sample preparation may not translate to other of murine faecal pellets (e.g. young vs. old individuals) or other mammalian faeces (e.g. *H. sapiens*) and would suggest a required sample preparation optimisation for each cohort. To conclude, both sample preparation approaches have advantages, and the choice may ultimately come down to which bacterial organism is under-investigation [25].

### Single-step database search allows optimal control of false discovery rate

Currently, many metaproteomic studies use two-step database searches as a way to boost identification rates [36]. However, we demonstrate that this type of search dramatically underrepresents the number of false positives, due to the use of a decoy search strategy that is unsuitable in this context. Our results elaborate on a previous study by Muth and co-workers, who also emphasised the drawbacks of using a two-step search together with decoy strategy [48]. Using a single human LC-MS/MS measurement, our findings were so extreme that the number of false positives was equal or greater to the number of false negatives, with FDR outside of any accepted range (i.e. factual FDR > 0.1).

Using metaproteome samples of known composition, we expanded our investigation of search strategies by including “two-step taxa” and “two-step two sections”. The “two-step two sections” approach, implemented according to Kumar and colleagues [37], provided a middle ground in performance between the “two-step protein” and single-step search strategies, but at the expanse of much longer processing time. Nonetheless, our results confirmed the inability of two-step searches to control the FDR, including in context of metaproteomic samples. We argue that the use of a two-step search should be avoided whenever possible and replaced by alternative strategies, such as taxonomic foreknowledge or using matching metagenomes [49].

### Accurate taxonomic annotation of murine faecal samples can be generated by the Unipept software

Previous studies have shown that it is possible to derive taxonomic representation from MS- identified peptides of known taxonomic origin [32,33,50]. However, to the best of our knowledge, there has not been a comparison of software for the taxonomic annotation of peptides with unknown origin. Here, we compared three software packages, namely Kraken2 [38], Diamond [39] and Unipept [28], which use different algorithms to perform such taxonomic annotation. Using metaproteome samples of known composition, as well as metaproteome samples from 38 murine faeces, we determined that the Unipept software provided superior performance (i.e. precision, sensitivity). Notably, Unipept is very user- friendly, fast and was designed to work on MS-identified peptides [28]. Whereas, Diamond and Kraken2 have both been designed to work on full protein/gene sequences or nucleotide sequencing reads (as opposed to peptides), which may have contributed to their lower performance [38, 39]. Our assessments (i.e. sensitivity, specificity) were based on exact taxonomic identity and ignored hits from closely related taxon, which may have negatively affected the performance estimates of Kraken2 [38]. While, Unipept was clearly the optimal taxonomic annotation software for MS-identified peptides, it is currently limited to UniProt proteins, NCBI taxonomic hierarchy and trypsin cleavage.

### The metaproteome shows an enrichment in functionally-active pathways compared to the matching metagenomic potential

Here, we observed an overall positive correlation between gene and protein abundances derived from metaproteome and matching-metagenome analysis. This was previously reported in a longitudinal study of metaproteome/metagenome fluctuations from one individual with Crohn’s Disease [51]. In our case the significantly correlated entries were associated with core bacterial metabolic functions, such as carbon and energy metabolism or electron transfer activity [52]. Despite such correlations, we also reported extensive differences in quantified functions between metagenomics and metaproteomics. Notably, with regard to genetic information processing (KEGG level 2), the ribosome pathway was over-represented in entries with higher abundance in metaproteomes, whereas pathways associated with DNA repair, replication or recombination were over-represented in entries with increased abundance in metagenomes. This greatly highlights the main advantage of metaproteomics, which captures functionally active pathways, as opposed to the genetic potential represented by metagenomics [53]. Thus, these approaches are complementary to each other and can provide a more comprehensive understanding of a biological system.

## Conclusion

To conclude, in this study we present an integrated analytical and bioinformatic workflow to improve protein identification, taxonomic and functional coverage of the murine faecal metaproteome. LSC combined with in-solution digestion provided the highest identification rates, although leading to a potential enrichment in specific taxa. We also show that fast and accurate MS data processing can be achieved using a single-step database search. Taxonomic annotation can be generated directly from MS-based peptide identification using the Unipept software. While protein and gene abundances displayed an overall positive correlation, the metaproteome showed a significant functional enrichment compared to its metagenomic potential; thus, emphasizing the need for more metaproteomic studies for adequate functional characterisation of the microbiome.

## Methods

### Animals and faecal samples collection

Mouse faecal pellets obtained from a small cohort of six wild-type B6EiC3SnF1/J mice were used to compare sample purification and protein extraction methodologies from faeces (**Figure S1A**). A larger cohort of 38 mice (euploid and trisomic Ts65Dn) was used to obtain mouse faeces, for further assessment of the data analysis workflow. Mice were housed and faeces were collected following the experimental procedures evaluated by the local Ethical Committee (Barcelona Biomedical Research Park, Spain). Faecal pellets were collected fresh, placed at - 20 °C and stored at -80 °C until analysis.

### DNA extraction and whole-genome sequencing

Whole genome analysis was performed on the mouse cohort used for data analysis assessment. In brief, DNA was extracted from faecal samples using the FastDNA SPIN Kit (MP Biochemicals) and following manufacturer’s instructions. DNA concentration was measured using a Qubit fluorometer (Invitrogen) and samples were shipped frozen to the Quantitative Biology Centre (QBiC) at the University of Tuebingen for whole genome sequencing. Sequence data were generated on an Illumina HiSeq 2500 instrument (chemistry SBS v3 plus ClusterKit cBot HS) and processed as described previously [54] but with minor modifications that follow. Supplied sequence data were checked using fastQC v0.11.5 [55]. Data were trimmed with Trim Galore! (--clip_R1 10 --clip_R2 10 --three_prime_clip_R1 10 -- three_prime_clip_R2 10 --length 50; Babraham Bioinformatics). Mouse DNA within samples was detected by mapping reads against the mouse genome (GRCm38). Mouse-filtered read files (with an average of 3.58 ± 0.08 Gb sequence data per sample) were used for all subsequent analyses. Kraken2 2.0.8-beta [38] with the pre-compiled Genome Taxonomy Database [56] Functional annotation was achieved by mapping centroid protein sequences generated as described before [38, 54] using the eggNOG-mapper software (v.1.0.3) [57] and associated database (v.4.5).

### Sample treatment, cell lysis and protein extraction

Mouse faecal pellets obtained from wild-type B6EiC3SnF1/J mice were used to compare sample initial preparation methodologies (**Figure S1A**).

For the LSC procedure, faeces (∼50 mg) were resuspended in phosphate buffer (50 mM Na2HPO4/NaH2PO4, pH 8.0, 0.1 % Tween 20, 35x volume per mg) by vortexing vigorously for 5 min using 4 mm glass beads (ColiRollers^TM^ Plating beads, Novagen), followed by incubation in a sonication bath for 10 min and shaking at 1,200 rpm for 10 min in a Thermomixer with a thermo block for reaction tubes. Insoluble material was removed by centrifugation at 200 × g at 4 °C for 15 min. The supernatant was removed and the remaining pellet was subjected to two additional rounds of microbial cell extraction. After merging supernatants, microbial cells were collected by centrifugation at 13,000 × g at 4 °C for 30 min. The pellet was resuspended in 80 µL sodium dodecyl sulfate (SDS) buffer (2 % SDS, 20 mM Tris, pH 7.5; namely pellet extraction buffer) and heated at 95 °C for 30 min in a Thermomixer. The resulting suspension was divided into two parts to obtain technical replicates for the rest of the sample preparation workflow. Protein extraction was performed by cell homogenization using 0.1 mm glass beads (100 mg, SartoriusTM Glass Beads) for each replicate and the FastPrep-24 5G instrument (MP) at 4 m/s or BeadBug microtube homogenizer (BeadBug) at 4,000 rpm. Three cycles of homogenization including 1 min bead beating, 30 sec incubation at 95 °C, and 30 sec centrifugation at 13,000 × g were performed. The homogenate was diluted with 800 µL MgCl2 buffer (0.1 mg/mL MgCl2, 50 mM Tris, pH 7.5) and centrifuged at 13,000 rpm for 15 min. Proteins from the supernatant were precipitated overnight in acetone and methanol at 20 °C (acetone:methanol:sample with 8:1:1 ratio). Protein pellets were resuspended in 120 µL denaturation buffer (6 M urea, 2 M thiourea, 10 mM Tris, pH 8.0) for downstream use.

For the nLSC procedure, mouse faeces (∼25 mg) were homogenised directly in 150 µL pellet extraction buffer as described above with the following changes. A bead mixture of 0.1 mm glass beads (100 mg), 5 × 1.4 mm ceramic beads (Biolab products), and 1 × 4 mm glass bead was used for five cycles of homogenisation to break-up the faecal material.

### Protein digestion

Following extraction, protein amount was quantified using Bradford assay (Bio-Rad, Munich, Germany) [58] and two methods were compared to digest proteins extracted from LSC or nLSC procedures.

The in-solution digestion method was performed as follows. Proteins (20 µg starting material) were reduced in 1 mM dithiothreitol (DTT) and alkylated in 5.5 mM iodoacetamide at room temperature (RT) for 1 h each. Proteins were pre-digested with LysC at RT for 3 h using a protein to protease ratio of 75:1. Samples were diluted nine-fold with 50 mM ammonium bicarbonate and digested overnight with trypsin (Sequencing Grade Modified Trypsin, Promega) at pH 8.0 using a protein to protease ratio of 75:1.

Filter-aided sample preparation (FASP) was performed as previously published [59]. Briefly, proteins (10 µg starting material) were reduced in 0.1 M DTT for 40 min at RT. The reduced samples were added to the filter units (30 kDa membrane cut off) and centrifuged at 14,000 × g for 15 min. All further centrifugation steps were performed similarly unless otherwise noted. Samples were then washed with 2X 200 µL urea buffer (100mM Tris/HCl, pH 8.5, 8M urea) and centrifuged. Proteins were incubated in 50 mM IAA for 20 min at RT in the dark. After alkylation, samples were centrifuged and washed three times with 100 µL urea buffer. This was followed by three wash steps with 50 mM ammonium bicarbonate (ABC) for 10 min. Proteins were digested overnight at 37 °C using trypsin digestion (Sequencing Grade Modified Trypsin, Promega) at pH 8.0 using a protein to protease ratio of 100:1. On the following day, the peptides were centrifuged into fresh tubes at 14,000 × g for 10 min. An additional 40 μL ABC buffer was added to the filter units and this solution was also centrifuged to increase the peptide yield. To stop the digestion from either in-solution or FASP workflows, the samples were acidified to pH 2.5 with formic acid and cleaned for LC-MS/MS measurement using Empore C18 disks in StageTips [60].

### LC-MS/MS measurements

Samples were measured on an EASY-nLC 1200 (Thermo Fisher Scientific) coupled to a Q Exactive HF mass spectrometer (Thermo Fisher Scientific). The samples prepared for the sample purification and protein extraction methodologies assessment were all measured in duplicates to assess instrument reproducibility. Peptides were chromatographically separated using 75 μm (ID), 20 cm packed in-house with reversed-phase ReproSil-Pur 120 C18-AQ 1.9 μm resin (Dr. Maisch GmbH).

Peptide samples generated as part of the laboratory method optimisation (LSC vs. nLSC, FASP vs. in-solution) were eluted over 43 min using a 10 to 33 % gradient of solvent B (80 % ACN in 0.1 % formic acid) followed by a washout procedure. Peptide samples generated as part of the data analysis assessment (metaproteome vs. metagenome) were eluted over 113 min using a 10 to 33 % gradient of solvent B (80 % ACN in 0.1 % formic acid) followed by a washout procedure.

MS1 spectra were acquired between 300-1,650 Thompson at a resolution of 60,000 with an AGC target of 3 × 10^6^ within 25 ms. Using a dynamic exclusion window of 30 sec, the top 12 most intense ions were selected for HCD fragmentation with an NCE of 27. MS2 spectra were acquired at a resolution of 30,000 and a minimum AGC of 4.5 × 10^3^ within 45 ms.

### LC-MS/MS data processing

Raw data obtained from the instrument were processed using MaxQuant (version 1.5.2.8) [61]. The protein sequence databases used for database search consisted of the complete *Mus musculus* Uniprot database (54,506 sequences) and frequently observed contaminants (248 entries), as well as the mouse microbiome catalogue (∼2.6 million proteins) [40] for the raw data from laboratory method optimisation samples or the matching metagenome gene translation (∼1.5 million proteins) for the raw data from data analysis assessment samples. A FDR of 1 % was required at the peptide and protein levels. A maximum of two missed cleavages was allowed and full tryptic enzyme specificity was required. Carbamidomethylation of cysteines was defined as fixed modification, while methionine oxidation and N-terminal acetylation were set as variable modifications. Match between runs was enabled where applicable. Quantification was performed using label-free quantification (LFQ) [62] and a minimum peptide count of 1. All other parameters were left to MaxQuant default settings.

### Comparison of sample preparation methods

Unless stated otherwise, the analyses described below were performed in the R environment [63]. To compare the different centrifugation, digestion and lysis methods, we counted for each sample the number of peptide and protein groups with intensities and LFQ intensities superior to zero, respectively. We tested for significant differences between methods using unpaired t- tests via the ggplot2 package [64]. Quantified peptides and protein groups were checked for overlap between the centrifugation methods using the VennDiagram package. The proportion of host (*Mus musculus*) proteins was computed by summing up all host proteins iBAQ values and then dividing by the total iBAQ per sample. The centrifugation methods were evaluated using an unpaired t-test.

The taxonomy representation, for the centrifugation methods, was done via the Unipept online software (v. 4.5.1) [34]. The quantified peptides (intensity superior to zero) were imported into Unipept with I-L not equal. The Unipept result were used to count the number of non-redundant peptides assigned to each taxonomic node.

For the differential protein abundance analysis (between LSC and nLSC), the MSnBase package was used as organisational framework for the protein groups LFQ data [65]. Host proteins, reverse hit and potential contaminant proteins were filtered out. Protein groups were retained for further analysis only if more than 90 % of samples within either LSC or nLSC group had an LFQ superior to the first quartile overall LFQ. Significantly changing proteins were identified using paired t-test. Significance was set at an adjusted p-value of 0.01 following Benjamini-Hochberg multiple correction testing, as well as a minimum LSC/nLSC fold-change of ±1.5. The over-representation and GSEA testing of KEGG pathways were done for the significantly up- and down-regulated proteins as well as for the proteins uniquely identified per group via the clusterProfiler package based on hypergeometric distribution (p-adj. ≤ 0.05) [66].

### Single- versus two-step search assessment using HeLa cell line sample

HeLa cells were prepared for LC-MS/MS measurements using published method [67]. Briefly, cells were grown in DMEM medium and harvested at 80 % confluence. Proteins were precipitated using acetone and methanol. Proteins were reduced with DTT and digested with Lys-C and trypsin. Peptides were purified on Sep-Pak C18 Cartridge.

Sample was measured as described in the LC-MS/MS measurements section but for a few changes. Peptide sample was eluted over 213 min using a 7 % (0 min), 15 % (140 min) and 33 % (213 min) gradient of solvent B (80 % ACN in 0.1 % formic acid) followed by a washout procedure. The top 10 most intense ions were selected for HCD fragmentation.

Raw data were processed as described in the LC-MS/MS data processing section with a few alterations. Match between runs was disabled. The protein sequence databases used for database search consisted of the complete *Homo sapiens* Uniprot database (93,799 sequences), frequently observed contaminants (248 entries), as well as the mouse microbiome catalogue (∼2.6 million proteins) [40]. Several processings were performed differing in the number of microbiome catalogue entries included, which led to an increase in database size of 0×, 1×, 2×, 5×, 10× and 20× compared to the *H. sapiens* database alone. These processings also differed in the database search strategies used, namely single- or two-step search [36].

Identified MS/MS, peptides and protein groups were assigned to kingdom of origin (conflicts were resolved to Eukaryota by default). To compare the different database search strategies, we counted the number of identified MS/MS, non-redundant peptides and protein groups associated to each kingdom (as well as reverse hits and potential contaminants). We also calculated the FDR based solely on reverse hits or together with bacterial hits (factual FDR) in order to investigate the true number of false positives.

### Database search strategies assessment using known microbiome samples

We used the samples generated by Kleiner and colleagues, specifically the uneven organisms preparation described in the earlier publication [32]. This dataset contained LC-MS/MS measurements (N = 8) that we processed as described in the LC-MS/MS data processing section with a few alterations. Match between runs was disabled. The protein sequence databases used for database search consisted of the proteome of all 32 organisms present in the synthetic samples (“uneven database” = 122,972 sequences), frequently observed contaminants (248 entries), as well as the mouse microbiome catalogue (∼2.6 million proteins) [40]. Several processings were performed differing in the number of microbiome catalogue entries included, which led to an increase in database size of 0×, 0.5×, 1×, 2×, 5×, 10× and 20× compared to the “uneven database” alone. These processings also differed in the database search strategies used, namely single-step search, “two-step protein” search to keep identified proteins [36], “two-step taxa” search to keep identified taxa [30], and “two-step two sections” search to keep identified proteins after sectioned search [37].

Identified protein groups were assigned to database of origin, namely “uneven database” or mouse microbiome catalogue database. For each sample, this allowed computation of the number of (1) true positive hits, must be hits from the “uneven database”; (2) false positive hits, must be hits from the mouse microbiome catalogue; (3) false negative hits, the total identified protein count in the “uneven database” (total from 8 samples) minus the true positives; and (4) true negative hits, the total protein count in the mouse microbiome catalogue minus the false positives. This allowed calculation of the accuracy, precision and sensitivity for each increase in the database size. We also calculated the factual FDR based on reverse hits together with mouse microbiome catalogue hits in order to investigate the true number of false positives.

Using only the processings against the largest database (20×), we filtered our data for protein groups with a minimum of one or two unique peptides. The true positive count and factual FDR were calculated (and compared) for each combination of search strategy and filtering, as described in the previous paragraph.

### Taxonomic representation of known microbiome samples

We also used the uneven samples generated by Kleiner and colleagues [32] to investigate the taxonomic representation derived from MS-identified peptides. The gold-standard processing was used, with single-step database search against the proteome of all 32 organisms present in the synthetic samples (“uneven database” = 122,972 sequences). MS-identified peptides were submitted to (1) Kraken2 (v. 2.1.1) [38], (2) Diamond (v. 2.0.9) [39], or (3) Unipept online (v. 4.5.1) [28] software for taxonomic assignments. The protein sequences from Uniprot (swissprot and trembl) were used as database for each software. The Diamond alignment was performed using sensitive and taxonomic classification mode. The Unipept online analysis was done via the metaproteome analysis function with I-L not equal. The Kraken2 k-mer analysis was carried out in translated mode using back-translated peptide sequences (back-translation done with EMBOSS backtranseq). For each software approach, the complete taxonomic lineage (NCBI) was retrieved per peptide and the lowest common ancestor was determined.

For each sample, we determined and computed the number of taxa that are (1) true positive hits, must be an identified taxon used for the preparation of the synthetic samples; (2) false positive hits, must be an identified taxon not used for the preparation of the synthetic samples; (3) false negative hits, the total number of taxa used for the preparation of the synthetic samples minus the true positives; and (4) true negative hits, the total number of taxa (with at least one Uniprot protein) minus the true and false positives. This allowed calculation of the accuracy, precision, specificity, sensitivity and F-measure for different PSM count thresholds. Taxa were then quantified per sample based on the different software approaches by summing the peptide intensities and then normalised to percentage of total peptide intensities. At each taxonomic level, the Spearman’s rank correlation was calculated between the expected taxon representation in the uneven samples and the taxa representation determined from each software.

To investigate the taxonomic identification in context of different database search strategies, we performed the taxonomic annotation via Unipept for all uneven data processings described in the above section. We then carried out all steps described in the previous paragraph in order to compute the F-measure per search strategy and database size.

### Taxonomic representation of faecal microbiome samples

All subsequent sections use the faecal samples from a 38 mice cohort. These were prepared via LSC and in-solution protein digestion, as described above. The resulting peptide mixtures were measured on a Q Exactive HF mass spectrometer and processed against the matching metagenome gene translation, as described above.

The MS-identified peptides in this dataset were taxonomically annotated with Kraken2, Diamond and Unipept, as described above. Taxa were quantified as described above (sum of peptide intensities). The Spearman’s rank correlation in taxon representation was calculated for each pairwise combination of samples within software.

For each sample, we determined and computed the number of species that are (1) true positive hits, must be an identified species reported in the mouse microbiome catalogue; (2) false positive hits, must be an identified species not reported in the mouse microbiome catalogue; and (3) false negative hits, the total number of species reported in the mouse microbiome catalogue minus the true positives. This allowed calculation of the precision, sensitivity and F-measure for each samples and annotation software.

### Metagenome to metaproteome correlation

All subsequent sections use the faecal samples from a 38 mice cohort. These were prepared via LSC and in-solution protein digestion, as described above. The resulting peptide mixtures were measured on a Q Exactive HF mass spectrometer and processed against the matching metagenome gene translation, as described above. For direct comparison between metagenome and metaproteome, the identified genes were collapsed into groups identical to protein groups composition from mass spectrometry. Each gene groups abundance was calculated as the highest gene abundance within that group. Each gene groups and corresponding protein groups abundances were correlated across samples using Spearman’s rank correlation from the stats package. Significance was set at an adjusted p-value of 0.05 following Benjamini-Hochberg multiple correction testing. The GSEA testing of KEGG pathways and Gene ontologies were performed via the clusterProfiler package based on hypergeometric distribution (p-adj. ≤ 0.05) [66] following z-scoring of Spearman rho estimate per KEGG orthologies.

### Functional KEGG categories representation

For each sample, the protein groups iBAQ values were summed per KEGG category (level 2) on the basis of KEGG orthology annotation. The same approach was also undertaken for gene count. The KEGG category abundance were normalised for differing number of KO entries per category and for variation between samples; this was done separately for metagenome and metaproteome. Differences in KEGG category abundance between metagenome and metaproteome were tested using paired t-tests from the stats package. Significance was set at an adjusted p-value of 0.01 following Benjamini-Hochberg multiple correction testing. Significantly changing KEGG categories were prioritised based on gene groups to protein groups correlation (see section Metagenome to metaproteome correlation), whereby the Wilcoxon rank-sum test was used to identify KEGG category containing KO entries whose correlation differ from overall distribution (adjusted p-value ≤ 0.05).

To investigate further these selected KEGG categories, the protein groups iBAQ and gene count were used as described in the previous paragraph to derive KO normalised abundance and t-test results. Using the KO entries from each selected KEGG categories, separate GSEA testing of KEGG pathways were performed via the clusterProfiler package based on hypergeometric distribution (p-adj. ≤ 0.05).

## Acknowledgments

MED, BM, XA and LD are grateful to the European Community 7^th^ Framework Program under Coordinated Action NEURON-ERANET (grant agreement 291840). BM was supported by grants from the Deutsche Forschungsgemeinschaft (German Research Foundation Cluster of Excellence EXC 2124). BM and NN acknowledge support by the High Performance and Cloud Computing Group at the Center for Data Processing of the University of Tübingen, the state of Baden-Wuerttemberg through bwHPC. The metagenomic work detailed herein used the computing resources of the UK MEDical BIOinformatics partnership – aggregation, integration, visualization and analysis of large, complex data (UK Med-Bio) – which was supported by the Medical Research Council (grant number MR/L01632X/1). XA acknowledge support from the MINECO, Spain (grant number PCIN-2014-105).

## Author contributions

LD, XA, MD and BM designed the study. XA and CG generated the mouse cohorts and collected the murine faecal material. LMG extracted the DNA from all faecal samples prior to metagenomic sequencing. VA, TG and ID prepared the murine faecal samples for proteomic measurement by mass spectrometry. LH processed the metagenomic data, generating the taxonomic and gene abundance outputs. NN processed the metaproteomic datasets and performed the proteogenomic integration. NN wrote the manuscript with the input from all authors.

## Data Access

The complete metaproteomic bioinformatic workflow is available online [68]. The mass spectrometry proteomic data have been deposited to the ProteomeXchange Consortium via the PRIDE [69] partner repository with the dataset identifiers PXD020695, PXD020738, PXD021928, PXD021932 and PXD027306. Trimmed whole genome sequence data with mouse reads removed have been deposited with GenBank, EMBL and DDBJ databases under the BioProject accession PRJNA473429.

## Notes

### Competing Interest Statement

The authors have declared no competing interest.

