## Supplementary information for "An Integrated Workflow for Enhanced Taxonomic and Functional Coverage of the Mouse Faecal Metaproteome"

[1] Proteome Center Tuebingen, University of Tuebingen, Germany; [2] Biomolecular Medicine Section, Division of systems Medicine, Department of Metabolism, Digestion and Reproduction, Imperial College London, Sir Alexander Fleming building, London SW7 2AZ, UK; [3] Department of Biosciences, Nottingham Trent University, UK; [4] Bellvitge Biomedical Research Institute, Spain; [5] Université Côte d'Azur, CNRS, Inserm, IPMC, France; [6] Neurophysiology Unit, University of Barcelona - IDIBAPS, Spain; [7] Genomic and Environmental Medicine, National Heart & Lung Institute, Faculty of Medicine, Imperial College London, London, SW3 6KY, United Kingdom. [8] European Genomic Institute for Diabetes, INSERM UMR 1283, CNRS UMR 8199, Institut Pasteur de Lille, Lille University Hospital, University of Lille, 59045 Lille, France.

### Figure S1

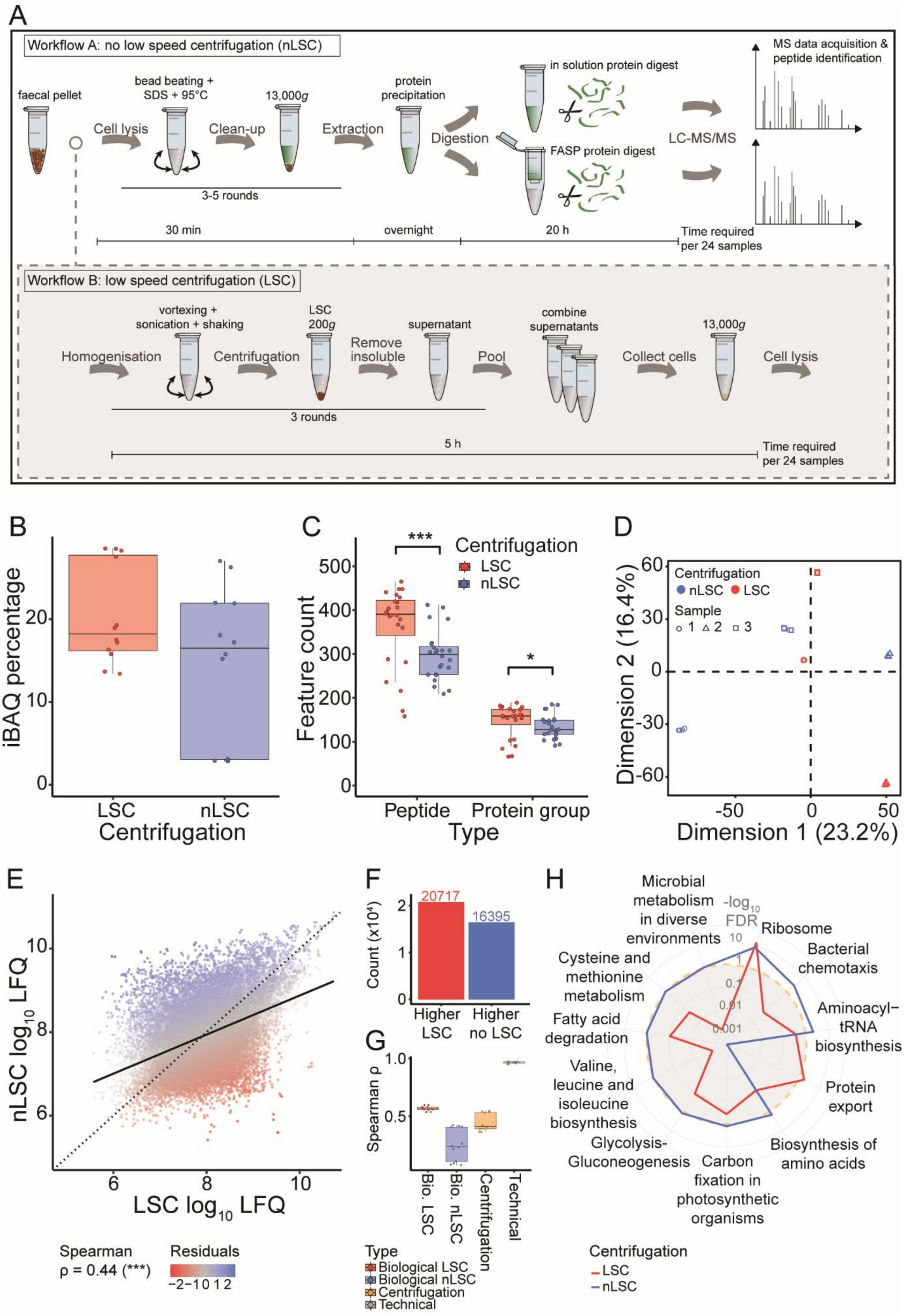

A
Workflow A: no low speed centrifugation (nLSC)
faecal pellet
bead beating + SDS + 95°C
13,000g
protein precipitation
in solution protein digest
MS data acquisition & peptide identification
Cell lysis
Clean-up
Extraction
Digestion
FASP protein digest
LC-MS/MS
3-5 rounds
30 min
overnight
20 h
Time required per 24 samples
Workflow B: low speed centrifugation (LSC)
vortexing + sonication + shaking
LSC 200g
supernatant
combine supernatants
13,000g
Homogenisation
Centrifugation
Remove insoluble
Pool
Collect cells
Cell lysis
3 rounds
5 h
Time required per 24 samples
B
iBAQ percentage
LSC
nLSC
Centrifugation
C
Feature count
Centrifugation
LSC
nLSC
Peptide
Protein group
Type
D
Centrifugation
nLSC LSC
Sample
1 2 3
Dimension 2 (16.4%)
Dimension 1 (23.2%)
E
nLSC log10 LFQ
LSC log10 LFQ
Spearman
ρ = 0.44 (***)
Residuals
F
Count (x10⁴)
20717
16395
Higher LSC
Higher no LSC
G
Spearman ρ
Bio. LSC
Bio. nLSC
Centrifugation
Technical
Type
Biological LSC
Biological nLSC
Centrifugation
Technical
H
-log10 FDR
Microbial metabolism in diverse environments
Ribosome
Bacterial chemotaxis
Aminoacyl−tRNA biosynthesis
Protein export
Biosynthesis of amino acids
Carbon fixation in photosynthetic organisms
Glycolysis-Gluconeogenesis
Valine, leucine and isoleucine biosynthesis
Fatty acid degradation
Cysteine and methionine metabolism
Centrifugation
LSC
nLSC

**Figure S1: Low speed centrifugation impacts protein identification and taxonomic representation.** A) Laboratory workflow overview for the nLSC and LSC approaches. For the LSC, faecal material is homogenised, centrifuged three times at 200 ×g to remove insoluble material and finally cells are collected by centrifugation at 13,000 ×g (accounting for an extra 5h per 24 samples). The LSC samples are subsequently prepared identically to the nLSC approach; i.e. cell lysis, protein extraction, protein digestion and LC-MS/MS measurement. An additional comparison focused on the protein digestion step either using Filter-Aided Sample Preparation (FASP) or in-solution. B) Percentage of the total iBAQ assigned to *Mus musculus* (host) protein groups per samples for the comparison between LSC (red) and nLSC (blue) methods. Represented significance results correspond to t-test on N = 12: * *p*- value ≤ 0.05, ** ≤ 0.01, *** ≤ 0.001. C) Number of identified peptides and protein groups per samples assigned to the host diet for the comparison between LSC (red) and nLSC (blue) methods. Represented significance results correspond to paired t-test on N = 24: * *p*- value ≤ 0.05, ** ≤ 0.01, *** ≤ 0.001. D) Principal component analysis showing separation of the samples based on biological replicates and sample preparation (LSC or nLSC) and the clustering of the technical replicates. E) Correlation in the protein groups LFQ values between the LSC and nLSC processed samples. F) This represents the count of protein groups with higher LFQ in LSC or nLSC approaches. G) This shows the protein groups LFQ correlation between LSC biological replicates (red), nLSC biological replicates (blue), centrifugation samples (orange) and technical MS-measurement replicates (grey). H) Radial plot of the over-represented KEGG pathways based on protein groups uniquely quantified in LSC (red) or nLSC (blue) processed samples. Significance threshold was set to an adjusted p-value ≤ 0.1.

### Figure S2

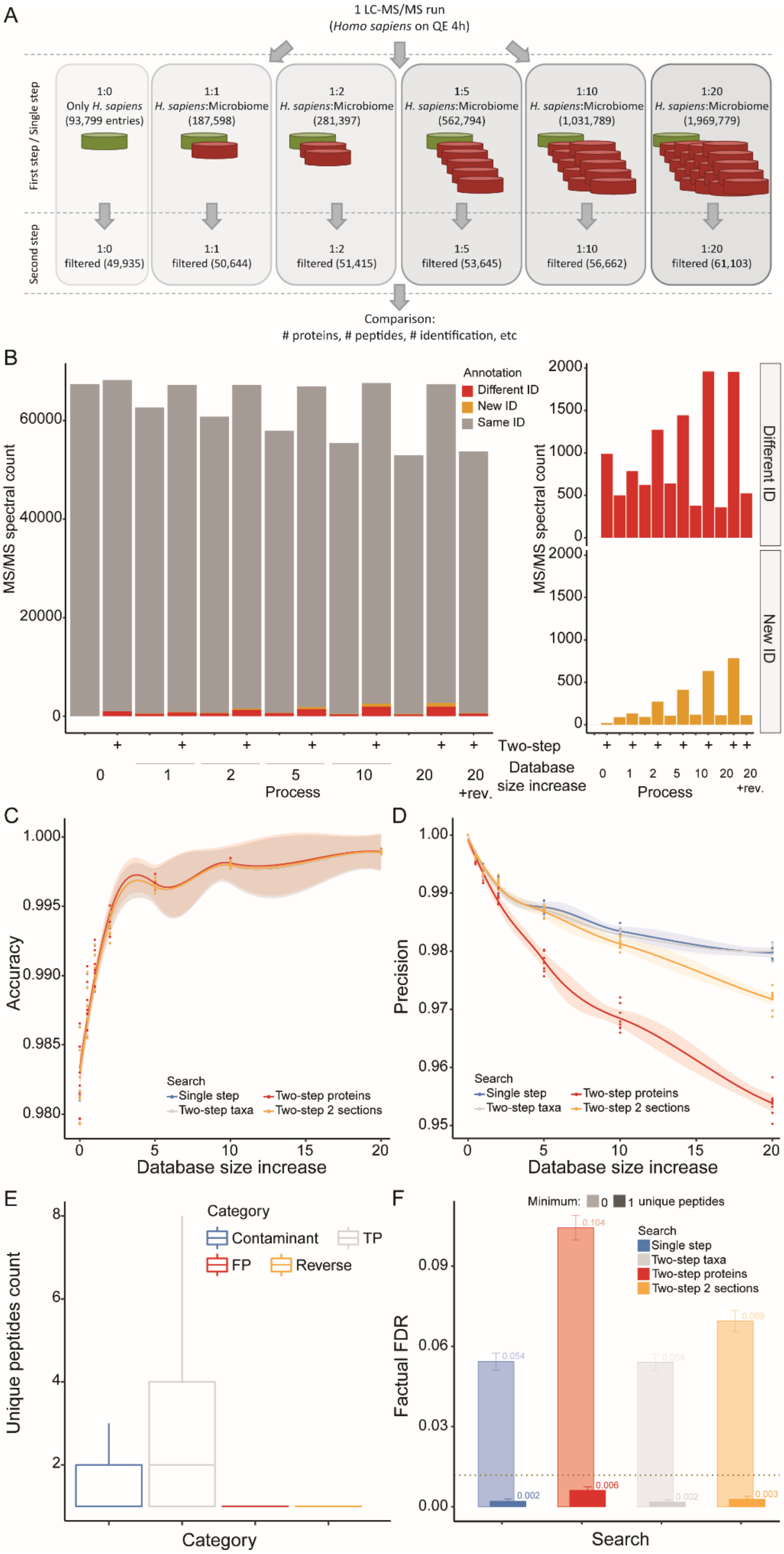

A
1 LC-MS/MS run
(Homo sapiens on QE 4h)
First step / Single step
Second step
1:0
Only H. sapiens
(93,799 entries)
1:1
H. sapiens:Microbiome
(187,598)
1:2
H. sapiens:Microbiome
(281,397)
1:5
H. sapiens:Microbiome
(562,794)
1:10
H. sapiens:Microbiome
(1,031,789)
1:20
H. sapiens:Microbiome
(1,969,779)
1:0
filtered (49,935)
1:1
filtered (50,644)
1:2
filtered (51,415)
1:5
filtered (53,645)
1:10
filtered (56,662)
1:20
filtered (61,103)
Comparison:
### proteins, # peptides, # identification, etc
B
MS/MS spectral count
Annotation
Different ID
New ID
Same ID
Two-step
Database size increase
Process
+rev.
Different ID
New ID
C
Accuracy
Search
Single step
Two-step taxa
Two-step proteins
Two-step 2 sections
Database size increase
D
Precision
Database size increase
E
Unique peptides count
Category
Contaminant
TP
FP
Reverse
F
Minimum:
0
1 unique peptides
Factual FDR
Single step
Two-step taxa
Two-step proteins
Two-step 2 sections
Search

**Figure S2: Two-step database search in combination with target-decoy strategy leads to a dramatic increase in false positive rate.** A) Bioinformatics workflow overview used to assess single- versus two-step search strategies. A single *Homo sapiens* LC-MS/MS run was searched against *H. sapiens* protein sequence database supplemented with increasing number of bacterial sequences (from 1:0, up to 1:20 human:bacterial sequences). B) The identified MS/MS spectra are counted for single- or two-step search strategies across increasingly large protein sequence databases. Colour coding corresponds to MS/MS spectra that were identified with identical sequence (grey), identified with different sequence (red) or newly identified (orange) in comparison to the processing against the human protein database only (gold standard). The two smaller panels (right) correspond to zoom on the MS/MS spectra identified with different sequence or newly identified. C & D) The accuracy (C) and precision (D) based on protein groups identification across increasingly large protein sequence databases. The compared database search strategies are single-step (blue), two-step taxon filtering (grey) and two-step protein filtering without (red) or with (orange) database sectioning. Lines represent the median (and the shading corresponds to the standard error) from N = 8 LC-MS/MS runs. E) Distribution of the number of unique peptides per protein groups is shown for a representative processing (i.e. single-step search against database size 20). Distribution is colour-coded per category, with true positive (grey), false positive (red), contaminant (blue) and reverse (orange) hits. F) The factual FDR based on protein groups identified with a minimum of one (shaded colouring) or two (unshaded colouring) unique peptides for the largest database (i.e. 20). The compared database search strategies are single-step (blue), two-step taxon filtering (grey) and two-step protein filtering without (red) or with (orange) database sectioning. Bars and numbers indicate the median FDR, while error bars correspond to the standard deviation, from N = 8 LC-MS/MS runs. The commonly used FDR threshold of 0.01 is indicated as a horizontal dotted line (gold).

### Figure S3

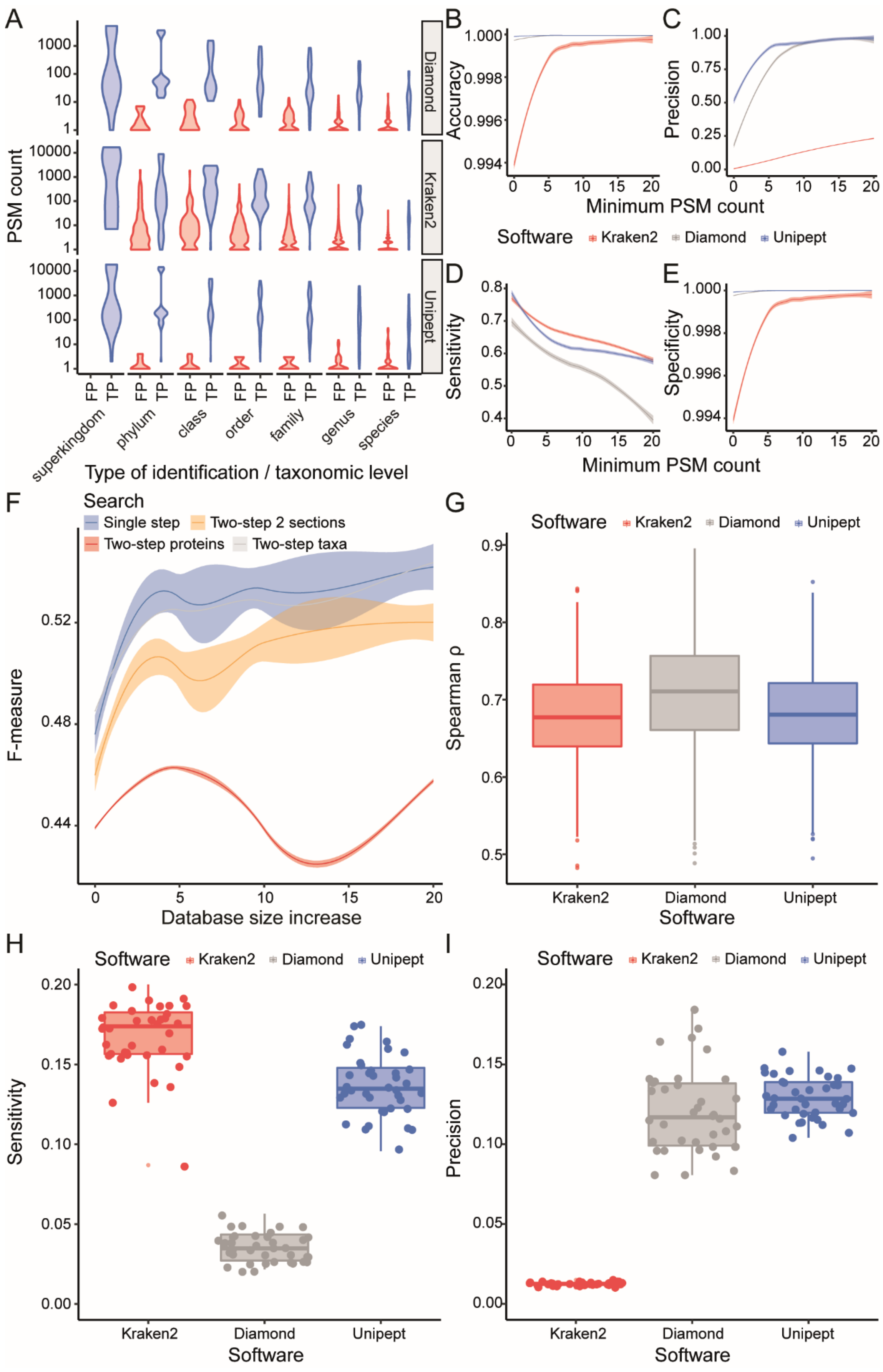

A
PSM count
1000
100
10
1
10000
Diamond
Kraken2
Unipept
FP
TP
superkingdom
phylum
class
order
family
genus
species
Type of identification / taxonomic level
B
Accuracy
1.000
0.998
0.996
0.994
C
Precision
1.00
0.75
0.50
0.25
0.00
Minimum PSM count
Software
Kraken2
Diamond
Unipept
D
Sensitivity
0.8
0.7
0.6
0.5
0.4
E
Specificity
Minimum PSM count
F
Search
Single step
Two-step 2 sections
Two-step proteins
Two-step taxa
F-measure
0.52
0.48
0.44
Database size increase
G
Spearman ρ
0.9
0.8
0.7
0.6
0.5
Software
H
Sensitivity
0.20
0.15
0.10
0.05
0.00
I
Precision

**Figure S3: Unipept software provides the most precise taxonomic annotation of MS-based peptide identification.** A) Distribution of PSM count is shown for false positive (red) and true positive (blue) taxon identification at each taxonomic level. The distribution is further split based on the software used for taxonomic annotation, i.e. Kraken2, Diamond and Unipept. B-E) Assessment of the impact of the minimum number of PSM count per taxon onto the accuracy (B), precision (C), sensitivity (D) and specificity (E) for taxonomic annotation. These quality metrics were compared between Kraken2 (red), Diamond (grey) and Unipept (blue) software. F) The F-measure based on taxon identification across increasingly large protein sequence databases. The compared database search strategies are single-step (blue), two-step taxon filtering (grey) and two-step protein filtering without (red) or with (orange) database sectioning. A-F) Samples from the study by Kleiner and colleagues, with N = 8. G) A comparison of the distribution in taxon abundance correlation (Spearman ρ) between different taxonomic annotation software (i.e. Kraken2, Diamond and Unipept). The correlation are computed between every combination of samples within each software. H-I) A comparison of the sensitivity (H) and precision (I) distribution for taxonomic annotation between the Kraken2 (red), Diamond (grey) and Unipept (blue) software. Each point represents an individual mouse. G-I) Samples from this study using mouse faecal material, with N = 38.

### Figure S4

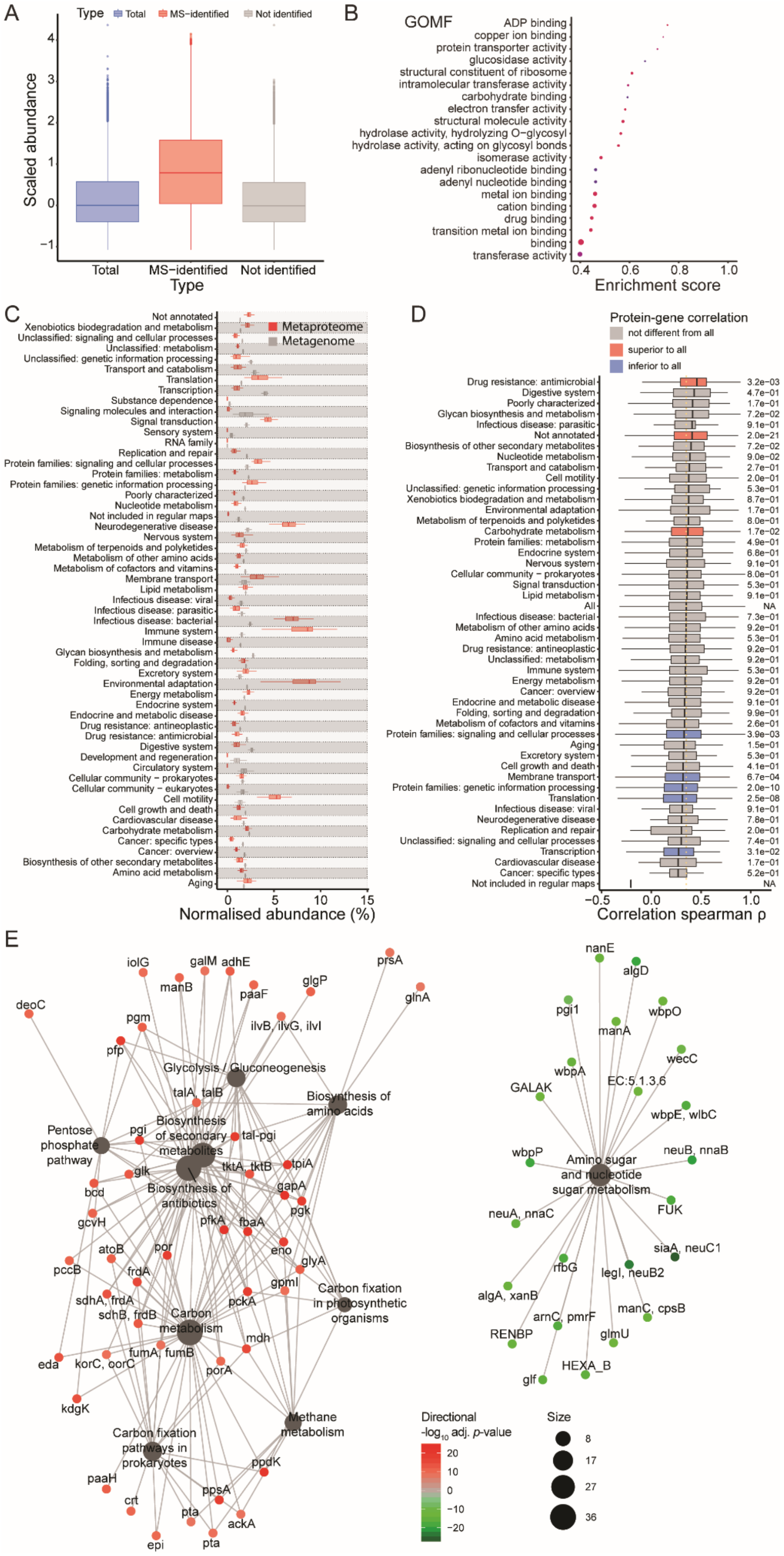
A
Type
Total
MS−identified
Not identified
Scaled abundance
Type
B
GOMF
ADP binding
copper ion binding
protein transporter activity
glucosidase activity
structural constituent of ribosome
intramolecular transferase activity
carbohydrate binding
electron transfer activity
structural molecule activity
hydrolase activity, hydrolyzing O−glycosyl
hydrolase activity, acting on glycosyl bonds
isomerase activity
adenyl ribonucleotide binding
adenyl nucleotide binding
metal ion binding
cation binding
drug binding
transition metal ion binding
binding
transferase activity
Enrichment score
C
Metaproteome
Metagenome
Not annotated
Xenobiotics biodegradation and metabolism
Unclassified: signaling and cellular processes
Unclassified: metabolism
Unclassified: genetic information processing
Transport and catabolism
Translation
Transcription
Substance dependence
Signaling molecules and interaction
Signal transduction
Sensory system
RNA family
Replication and repair
Protein families: signaling and cellular processes
Protein families: metabolism
Protein families: genetic information processing
Poorly characterized
Nucleotide metabolism
Not included in regular maps
Neurodegenerative disease
Nervous system
Metabolism of terpenoids and polyketides
Metabolism of other amino acids
Metabolism of cofactors and vitamins
Membrane transport
Lipid metabolism
Infectious disease: viral
Infectious disease: parasitic
Infectious disease: bacterial
Immune system
Immune disease
Glycan biosynthesis and metabolism
Folding, sorting and degradation
Excretory system
Environmental adaptation
Energy metabolism
Endocrine system
Endocrine and metabolic disease
Drug resistance: antineoplastic
Drug resistance: antimicrobial
Digestive system
Development and regeneration
Circulatory system
Cellular community − prokaryotes
Cellular community − eukaryotes
Cell motility
Cell growth and death
Cardiovascular disease
Carbohydrate metabolism
Cancer: specific types
Cancer: overview
Biosynthesis of other secondary metabolites
Amino acid metabolism
Aging
Normalised abundance (%)
D
Protein-gene correlation
not different from all
superior to all
inferior to all
Drug resistance: antimicrobial 3.2e−03
Digestive system 4.7e−01
Poorly characterized 1.7e−01
Glycan biosynthesis and metabolism 7.2e−02
Infectious disease: parasitic 9.1e−01
Not annotated 2.0e−21
Biosynthesis of other secondary metabolites 7.2e−02
Nucleotide metabolism 9.0e−02
Transport and catabolism 2.7e−01
Cell motility 2.0e−01
Unclassified: genetic information processing 5.3e−01
Xenobiotics biodegradation and metabolism 8.7e−01
Environmental adaptation 1.7e−01
Metabolism of terpenoids and polyketides 8.0e−01
Carbohydrate metabolism 1.7e−02
Protein families: metabolism 4.9e−01
Endocrine system 6.8e−01
Nervous system 9.1e−01
Cellular community − prokaryotes 8.0e−01
Signal transduction 5.3e−01
Lipid metabolism 9.1e−01
All NA
Infectious disease: bacterial 7.3e−01
Metabolism of other amino acids 9.2e−01
Amino acid metabolism 5.3e−01
Drug resistance: antineoplastic 9.2e−01
Unclassified: metabolism 9.2e−01
Immune system 5.3e−01
Energy metabolism 9.2e−01
Cancer: overview 9.2e−01
Endocrine and metabolic disease 9.1e−01
Folding, sorting and degradation 9.9e−01
Metabolism of cofactors and vitamins 2.6e−01
Protein families: signaling and cellular processes 3.9e−03
Aging 1.5e−01
Excretory system 5.3e−01
Cell growth and death 4.1e−01
Membrane transport 6.7e−04
Protein families: genetic information processing 2.0e−10
Translation 2.5e−08
Infectious disease: viral 9.1e−01
Neurodegenerative disease 7.8e−01
Replication and repair 2.0e−01
Unclassified: signaling and cellular processes 7.4e−01
Transcription 3.1e−02
Cardiovascular disease 1.7e−01
Cancer: specific types 5.2e−01
Not included in regular maps NA
Correlation spearman ρ
E
Glycolysis / Gluconeogenesis
Biosynthesis of amino acids
Pentose phosphate pathway
Biosynthesis of secondary metabolites
Biosynthesis of antibiotics
Carbon metabolism
Carbon fixation in photosynthetic organisms
Methane metabolism
Carbon fixation pathways in prokaryotes
Amino sugar and nucleotide sugar metabolism
Directional
-log10 adj. p-value
Size

**Figure S4: Functionally active pathways derived from the metaproteome differs from the metagenome potential.** A) Comparison of the distribution of gene abundances (median scaled to 0) based on all identified genes (blue) and genes whose protein was identified by mass spectrometry (red) or not (grey). B) Gene set enrichment analysis based on ranking of the protein/gene groups correlation for molecular function gene ontology categories. Category node colour corresponds to GSEA results adjusted p-value and node size matches the number of protein/gene group assigned to the category. C) Comparison in the proportion of all KEGG functional categories (level 2) between metaproteome (red) and metagenome (grey). Represented significance results correspond to paired t-test on N = 38. D) Protein groups to gene "groups" abundance correlation split per KEGG category (level 2). Categories are characterised by significantly superior (red) or inferior (blue) correlation in comparison to the overall distribution. Represented significance correspond to Wilcoxon signed-rank test at FDR $\leq$ 0.05. E) Interaction network between KEGG orthologies and KEGG pathways for the KEGG functional category "Carbohydrate metabolism". Pathway node size corresponds to number of KEGG orthologies associated to it. KEGG orthologies are colour-coded based on directional adjusted *p*-value from the t-test comparison between metaproteome and metagenome.
